# PIWIL3 forms a complex with TDRKH in mammalian oocytes

**DOI:** 10.1101/719963

**Authors:** Minjie Tan, Helena T.A. van Tol, David Rosenkranz, Elke F. Roovers, Mirjam J. Damen, Tom A.E. Stout, Wei Wu, Bernard A.J. Roelen

**Affiliations:** Department of Farm Animal Health, Faculty of Veterinary Medicine, Utrecht University, Utrecht, the Netherlands; Johannes Gutenberg-University Mainz, Institute of Organismic and Molecular Evolution, Anselm-Franz-von-Bentzel-Weg 7, 55128 Mainz, Germany; Biology of Non-coding RNA Group, Institute of Molecular Biology (IMB), Ackermannweg 4, 55128 Mainz, Germany; Biomolecular Mass Spectrometry and Proteomics, Bijvoet Center for Biomolecular Research and Utrecht Institute for Pharmaceutical Sciences, Utrecht University, Utrecht, the Netherlands; Netherlands Proteomics Centre, Utrecht, the Netherlands; Department of Equine Sciences, Faculty of Veterinary Medicine, Utrecht University, Utrecht, the Netherlands

## Abstract

PIWIs are crucial guardians of genome integrity, particularly in germ cells. While mammalian PIWIs have been primarily studied in mouse and rat, a homologue for the human PIWIL3 gene is absent in the Muridae family, and hence the unique function of PIWIL3 in germ cells cannot be effectively modeled by mouse knockouts. Herein, we investigated the expression, distribution and interaction of PIWIL3 in bovine oocytes. We localized PIWIL3 to mitochondria, and demonstrated that PIWIL3 expression is stringently controlled both spatially and temporally before and after fertilization. Moreover, we identified PIWIL3 in a mitochondrials-recruited three-membered complex with TDRKH and PNLDC1, and demonstrated by mutagenesis that PIWIL3 N-terminal arginine modifications are required for complex assembly. Finally we sequenced the piRNAs bound to PIWIL3-TDRKH-PNLDC1 and report here that about 50% of these piRNAs map to transposable elements, recapitulating the important role of PIWIL3 in maintaining genome integrity in mammalian oocytes.

## Introduction

Genomic integrity is critical for faithful propagation within the species. Ensuring genome integrity entails controlling transposable elements (TEs), which are mobile DNA sequences that can migrate within the genome^1^. Amongst various TEs, retrotransposons are especially damaging as these can be transcribed, reverse transcribed, and reinserted at multiple locations in the genome. While retrotransposon activity may be minimized by repressive DNA (cytosine) methylation, germ cells and early embryos are particularly vulnerable to retrotransposon reinsertion because in these developmental stages, cells undergo genome-wide demethylation^2^. Maintenance of genome integrity in oocytes and early developing embryos against TEs is therefore extremely important, since any genetic change would be passed on to the next generation.

In germ cells, P-element induced wimpy testis (PIWI) proteins that interact with 21-31 nucleotides non-coding PIWI interacting (pi)RNAs form an efficient system to silence TE activity^3–6^. The PIWI-piRNA pathway has been shown to suppress transposon activity both post-transcriptionally and transcriptionally^3,7^. Interestingly, PIWI-piRNA pathway activation has been mainly reported in the contexts of gametogenesis and embryology, further highlighting probably a unique and critical role for PIWIs in controlling genome stability, germ cell maturation and early embryonic development^8,9^.

Genetic mutation of different Piwi genes in mice (*Mili, Miwi*, and *Miwi2*) are associated with male sterility, but no abnormal phenotypes were observed in females^10–12^. This appears to imply that products of Piwi genes (homologues of human *PIWIL1, PIWIL2, PIWIL4*) are not critically needed during oogenesis, maturation and post-fertilisation, at least in the murine system. We have previously established that human, macaque and bovine oocytes express large amounts of piRNAs, and 30-50% of these are enriched for transposon sequences, in particular, the expression of a fourth PIWI gene, *PIWIL3*, was detected in human and bovine species but absent in murine systems^13,14^. Due, in part, to the perennial shortage of PIWIL3-specific antibodies, and the lack of mammalian species expressing PIWIL3 that are amenable to genetic manipulations, little is known about the function and subcellular localization of PIWIL3 as well as biogenesis in PIWIL3 related piRNAs.

Here we investigated the expression, distribution, and interaction of PIWIL3 in bovine oocytes by combining microinjection and immune-labeling techniques with affinity purification, crosslinking and mass spectrometry, as well as small RNA sequencing. We demonstrate here a novel and developmental stage-specific PIWIL3/TDRKH/PNLDC1 interaction network, which sheds light on piRNA biogenesis and maintenance of genome integrity in mammalian oocytes prior to and shortly after fertilization.

## Results

### PIWIL3 is expressed in the cytoplasm during oocyte maturation and early embryo development

Since the subcellular localization of a protein often dictates the mode of action and accessibility to substrates and cofactors, we sought to trace the intracellular distribution of PIWIL3 over developmental stages, to further elucidate the functional role of PIWIL3. While the intracellular distributions of PIWIL1, PIWIL2 and PIWIL4 have been elucidated in mouse testis^10,15^ and human fetal oocytes^14^, the subcellular compartment(s) in which PIWIL3 exerts its function remains uninvestigated to date. Herein, we follow the distribution and expression pattern of PIWIL3 in bovine oocytes and preimplantation embryos, by transiently expressing EGFP-PIWIL3 fusion proteins.

We generated EGFP-PIWIL3 fusion constructs (both N- and C-terminal tagging), performed *in vitro* transcription and directly injected *EGFP-PIWIL3* mRNA into bovine oocytes at the germinal vesicle (GV) stage and 2 pro-nuclei (2PN) stage zygotes after *in vitro* fertilization. Zygotes were cultured further *in vitro* for up to 8 days (blastocyst stage), during which embryos at various developmental stages post fertilization were harvested.

First we confirmed that the entire constructs were translated in the oocytes. Microinjection of the *EGFP-PIWIL3* and *PIWIL3-EGFP* fusion construct into oocytes resulted in proteins of 130 kDa, corresponding to the total length of EGFP (26 kDa) and PIWIL3 (100 kDa) (Figure 1a). Next, we followed the localization of EGFP-PIWIL3 in bovine oocytes. Starting from the GV stage, EGFP-PIWIL3 was localized largely to the oocyte cytoplasm in a punctate pattern but was excluded from the nucleus, whereas the EGFP control signal was distributed evenly throughout the cell. The expression of PIWIL3 remained cytoplasmic throughout preimplantation development but strongly reduced at the blastocyst stage (Figure 1b). The same localization patterns were observed using C-terminal EGFP tagging (Figure 1b). Collectively, these data support a largely cytoplasmic localization of PIWIL3 in bovine oocytes and preimplantation embryos.

**Figure 1:**
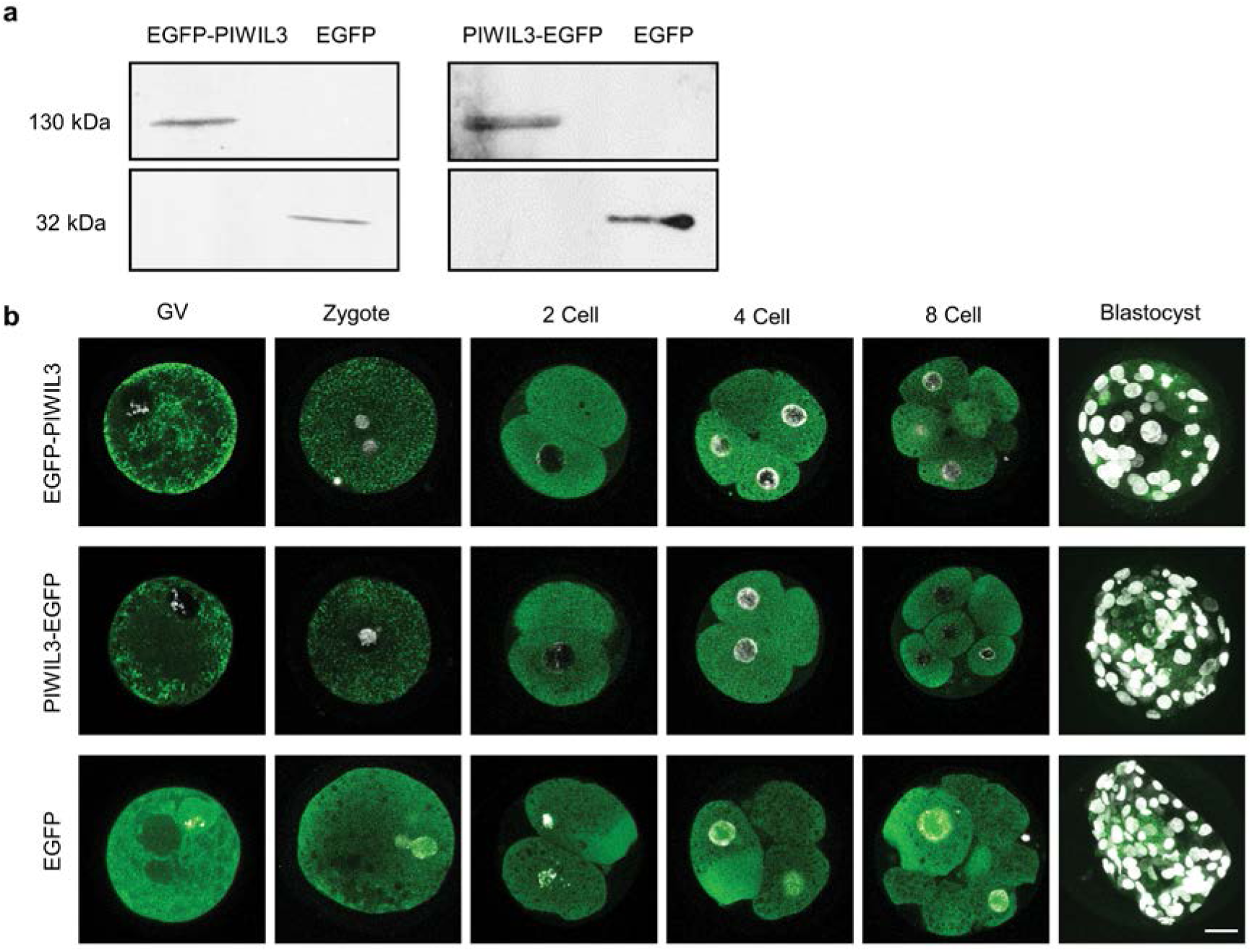
PIWIL3 is a cytoplasmic protein in oocytes and early stage embryos. a Detection of PIWIL3-EGFP, EGFP-PIWIL3 and EGFP in oocytes after mRNA injection by Western blotting. b Detection of EGFP and EGFP tagged PIWIL3 in oocytes and embryos by confocal laser scanning microscopy. Stages of oocyte and embryos are indicated at the top. DNA was stained with DAPI (white). Scale bar, 30 μm.

### PIWIL3 co-localizes with TDRKH on mitochondria

In the absence of a working PIWIL3 antibody, the functional complexes involving PIWIL3 have not been characterized. Here, we utilize an alternative approach to investigate the promising PIWIL3 binding partner in bovine oocytes, as guided by existing knowledge of other PIWI interactions.

Tudor domain containing proteins have been shown to interact with other PIWI proteins^16–18^, and we had established previously that Tudor and KH containing protein (TDRKH) is highly expressed in GV and Metaphase II (MII) oocytes^13^. We thus hypothesized that TDRKH might be a PIWIL3-associating protein in bovine oocytes. In further support of this possibility, we examined the temporal expression pattern of TDRKH in bovine gonads, and TDRKH levels in different stages of bovine oocytes. Using immunoblotting, high levels of TDRKH were detected in oocytes and testis while lower levels were detected in complete ovaries, indicating that TDRKH was specifically expressed in the germ cells and not in the somatic tissue of the gonads (Figure 2a). This was further illustrated by immunohistochemical analyses of testis and ovary sections which showed again the expression of TDRKH in germ cells but not somatic cells. In bovine testis, TDRKH expression was localized to immature and mature sperm cells (Figure 2b). In bovine ovaries, TDRKH expression was detected in oocytes, but not in granulosa and theca cells, at all stages from primary to antral follicles (Figure 2c).

**Figure 2:**
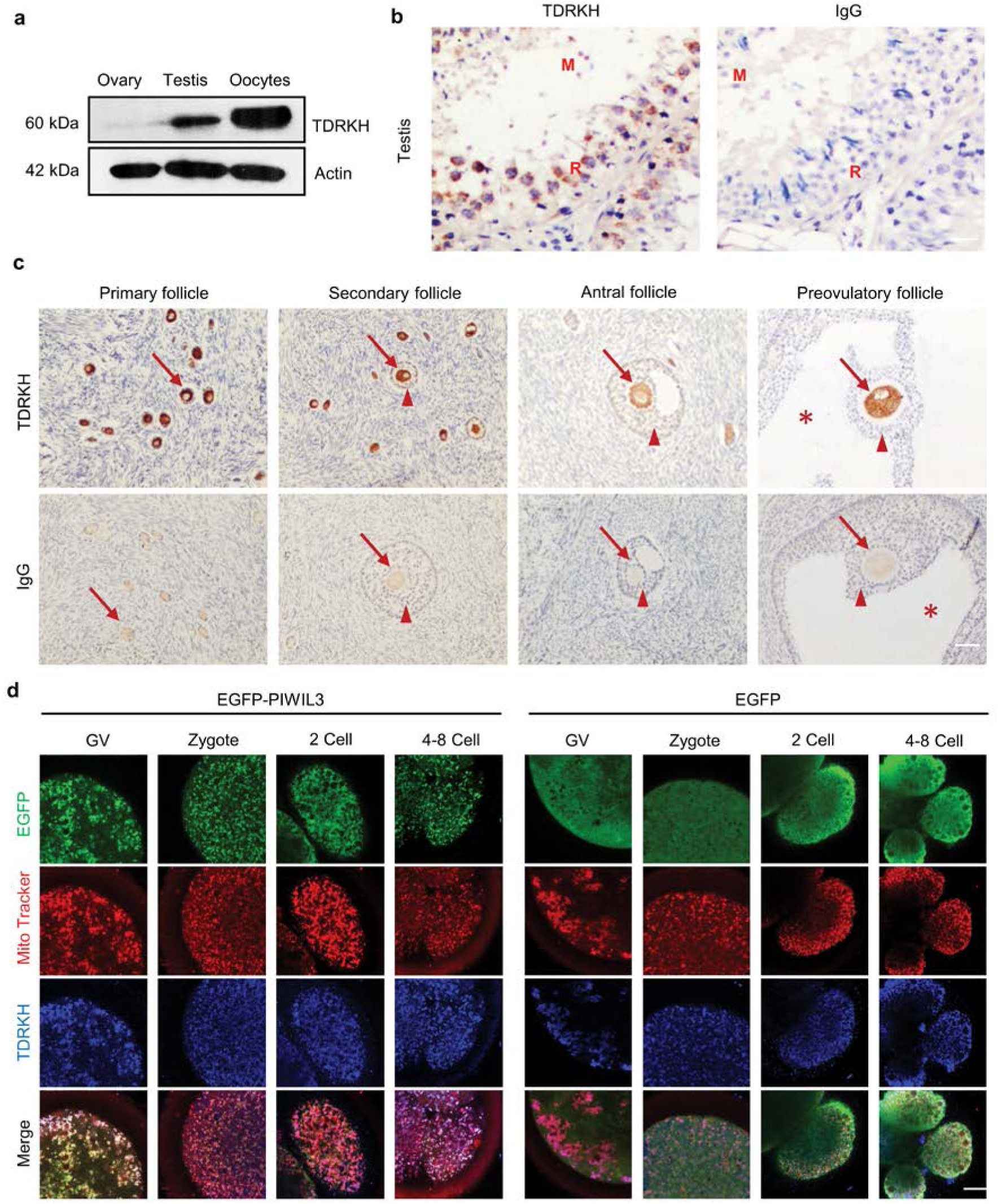
PIWIL3 colocalizes with TDRKH on mitochondria. **a** Western detection of TDRKH in bovine ovary, testis, as well as oocytes from antral follicles. **b** Immunostaining of TDRKH (brown staining) in paraffin sections of bovine testis. Blue: hematoxylin counterstaining. R: Round spermatid. M: Mature sperm. Scale bar, 100 μm. **c** Immunostaining of TDRKH (brown staining) in paraffin sections of bovine ovaries in different follicular stages. Blue: hematoxylin counterstaining. Arrows indicate oocytes; Triangles indicate cumulus cells; Asterisks indicate antrum. Scale bar, 100 μm. **d** Fluorescent detection of EGFP-PIWIL3 (green), EGFP (green), MitoTracker (mitochondria, red) and TDRKH (blue) in microinjected oocytes and early stage embryos. Scale bar, 20 μm.

Using the same EGFP fusion constructs described above and injection of *EGFP-PIWIL3* mRNA into bovine oocytes at the GV stage, we detected the co-localization of TDRKH with EGFP-PIWIL3 (and PIWIL3-EGFP) *in vivo*, in cytoplasmic regions that co-stain with MitoTracker (Figure 2d, Supplementary Figure S1). Collectively these data indicate that PIWIL3 and TDRKH co-localize at mitochondria.

### PIWIL3 is present in a complex with TDRKH and PNLDC1 in bovine oocytes

Since TDRKH and PIWIL3 fusion constructs co-localize in bovine oocytes, we were interested if these proteins also associate with each other.

We therefore proceeded to verify this interaction by reverse co-immunoprecipitation (co-IP) with a TDRKH antibody, followed by MS confirmation of the constituents in TDRKH complexes. To negate the possibility of non-specific proteins binding to GFP instead, we harvested bovine oocytes without mRNA injection for this experiment. By in-solution digestion of TDRKH co-IP eluates, the top three proteins detected were TDRKH, PNLDC1 and PIWIL3, making PIWIL3 and PNLDC1 highly confident interactors of TDRKH in bovine oocytes that were identified by many peptide spectra matches (Figure 3a, Supplementary Table S1). Other dominant proteins (OGDHL, IMMT, ARMC10 and PHB2) that showed enrichments in the immuno-purified TDRKH complexes were mostly mitochondrial, suggesting again that TDRKH connecting with PIWIL3 is docked on the mitochondria *in vivo*, in agreement with MitoTracker co-staining (Figure 2d and Supplementary Figure S1).

**Figure 3:**
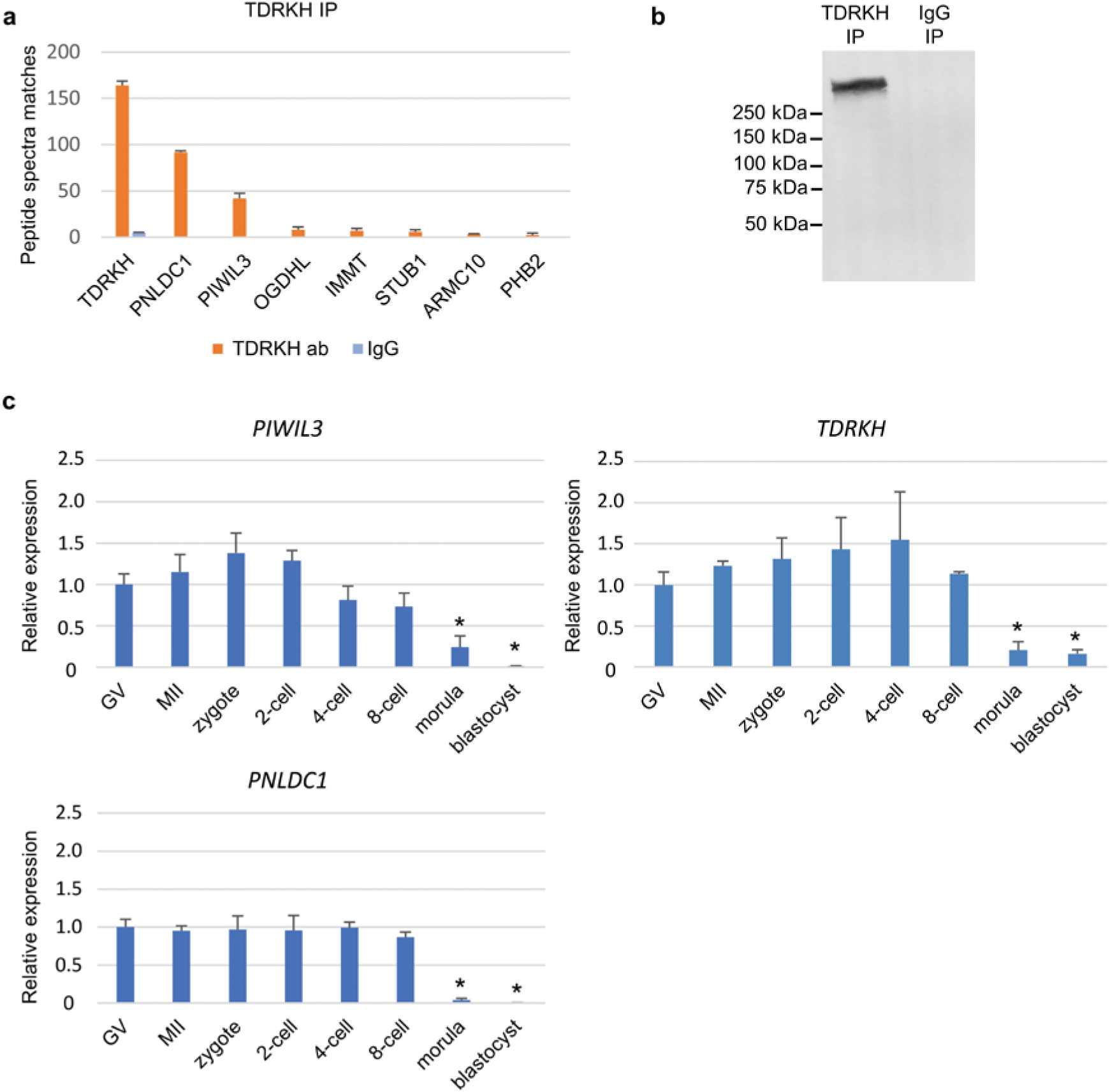
PIWIL3 forms a complex with TDRKH and PNLDC1 in oocytes. **a** Identification of TDRKH-associated proteins by mass spectrometry. Orange: TDRKH IP. Blue: IgG IP (negative control). **b** Crosslinked TDRKH complexes. Chemically crosslinked TDRKH complexes detected as a >250 kDa band by western blotting. **c** Bar graph showing *PIWIL3, TDRKH* and *PNLDC1* mRNA expression as detected by qRT-PCR in oocytes and early stage embryos. * p<0.05.

To further elucidate if PIWIL3 forms a complex with TDRKH and PNLDC1 simultaneously, or as two binary complexes (TDRKH-PIWIL3 and TDRKH-PNLDC1 respectively), we mildly crosslinked the proteins immuno-purified by TDRKH antibodies, eluted the complexes off Protein A/G-agarose beads, and detected by Western blotting a single complex species of >250kDa (Figure 3b). Since we did not observe two species of protein complexes containing TDRKH, and the theoretical combined size of PIWIL3-TDRKH-PNLDC1 alone is around 220kDa, we suspect that more proteins contribute to the complex. We report here that PIWIL3 is engaged in a TDRKH complex also including PNLDC1 in bovine oocytes.

Using qRT-PCR we examined the expression of *PIWIL3, TDRKH* and *PNLDC1* mRNA in oocytes and preimplantation embryos. The temporal expression pattern of these genes was similar, with expression in oocytes and embryos up to the 8-cell stage and decreased levels at morula and blastocyst stages (Figure 3c). Taken together, these results suggest that PIWIL3, TDRKH and PNLDC1 are collectively engaged in a germ cell specific function *in vivo*.

### PIWIL3 interacts with TDRKH through N-terminal arginines

To further characterize the mechanism of PIWIL3-TDRKH interactions, we generated PIWIL3-EGFP N-terminal deletion and substitution mutants (Figure 4a) and injected in vitro synthesized mRNA into GV stage oocytes, to examine the regions critical for TDRKH interaction. In other animal model systems, PIWI family proteins have been suggested to interact with Tudor domain-containing proteins through symmetrically dimethylated arginines at the N-terminus^16,17,19^. We demonstrate here that N-terminal deletion of the GRARVHARG motif from PIWIL3 (G3-G11) abolished PIWIL3 interaction with TDRKH and recruitment to the mitochondrial surface. This is exemplified by the diffuse pattern of N-terminal truncated PIWIL3-EGFP cytoplasmic mislocalisation (Figure 4b, De1 panels), confirming that the N-terminal GRARVHARG motif in PIWIL3 is indeed essential for TDRKH binding in a higher mammalian model system.

**Figure 4:**
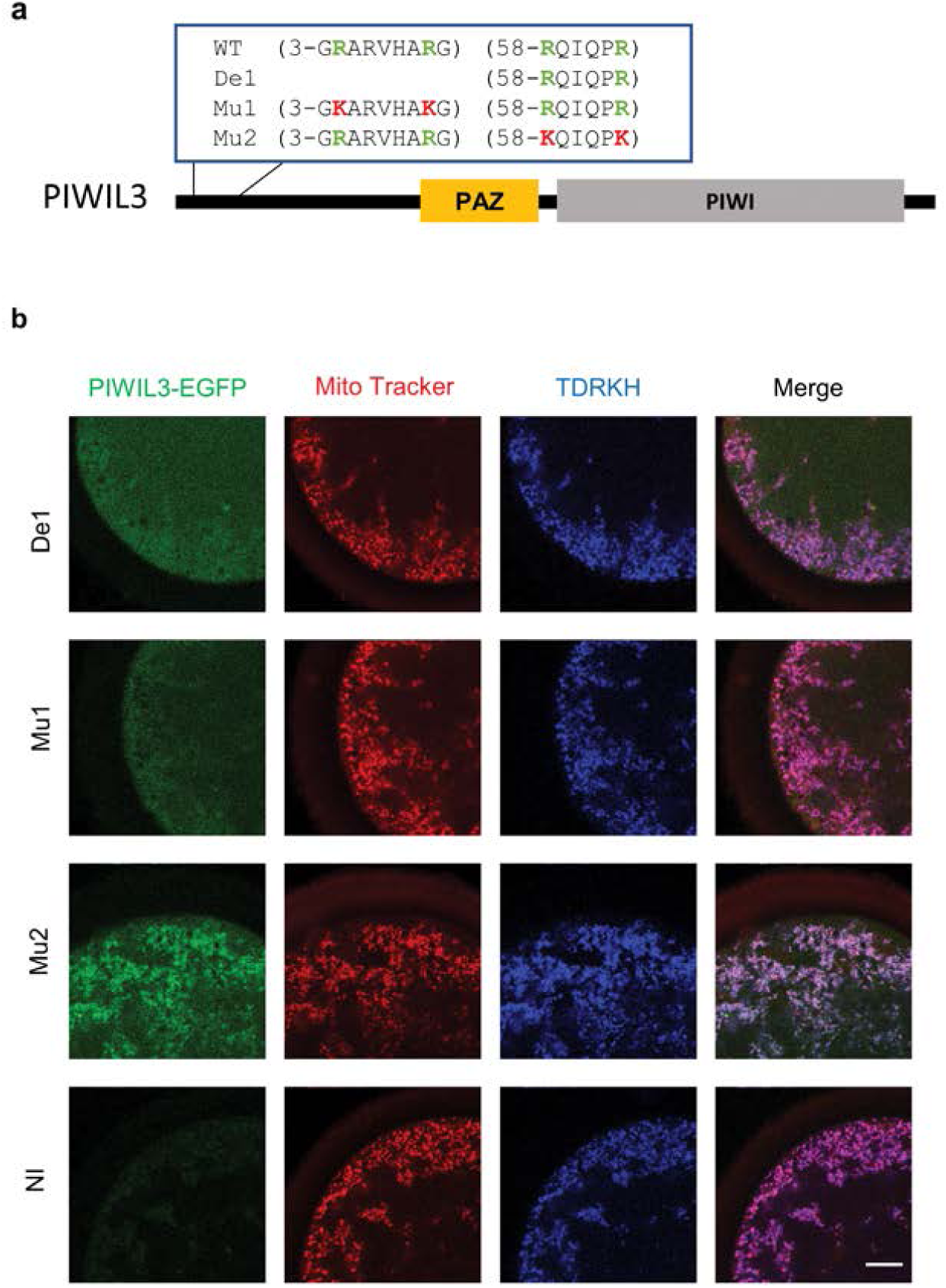
The N-terminal domain of PIWIL3 is critical for binding with TDRKH. **a** Domain map of PIWIL3 mutation construct. WT: wild type PIWIL3 sequence. De1: deletion construct lacking N-terminal GRARVHARG. Mu1/Mu2: mutation vectors. Green: original arginines; red: R→K substitutions. **b** Triple labeling of PIWIL3-EGFP mutants (green), mitochondria (red) and TDRKH (blue) in oocytes. Coding on the left indicates injected mRNAs. NI: Non-injected oocytes. Scale bar, 20 μm.

We next sought extensively for spectral evidence of symmetric dimethylations that might exist on these arginine residues in the G<u>R</u>ARVHA<u>R</u>G motif, by affinity purification and mass spectrometry of peptides derived from TDRKH-bound PIWIL3. This was however hampered by general difficulties in N-terminal sequence coverage in mass spectrometry. Thus, we turned instead to site-directed mutagenesis for alternative evidence that indicate the importance of arginine modifications in this N-terminal stretch. PIWIL3 arginine residues in this G<u>R</u>ARVHA<u>R</u>G motif (R4, R10) are likely critical to mediate TDRKH docking, as conservative substitutions of both arginines to lysines in full-length PIWIL3-EGFP was already sufficient to prevent TDRKH binding (Figure 4b, Mu1 panel). In contrast, conservative substitutions of R58 and R63, two dimethylated arginines detected by mass spectrometry on TDRKH-bound PIWIL3, had no functional impact on TDRKH binding and mitochondrial recruitment, serving as very convincing negative controls for this experiment. Hence, by a combination of mass spectrometry and N-terminal truncation/conservative amino acid substitutions, we provide evidence here that PIWIL3 interaction with TDRKH likely requires arginine modifications in the N-terminal G<u>R</u>ARVHA<u>R</u>G motif, in strong agreement with previous postulates in other species^16,17,19^.

We also compared the N terminal domains of PIWIL3 in different mammalian species: cow, human, rabbit, rhesus macaque and golden hamster together with mouse PIWIL4 (MIWI2) (Figure S4). A conserved GRAR region is present in the PIWIL3 N terminal region of these species. R4 and R10 are conserved in most of the species except in rabbit and golden hamster where R10 is replaced by respectively P and Q.

### piRNAs are loaded onto PIWIL3/TDRKH/PNLDC1 complexes

With our experimental evidence for the interaction between PIWIL3, TDRKH and PNLDC1, a new avenue to study PIWIL3-bound piRNA species arose, even in the absence of PIWIL3-specific antibodies. Instead of directly retrieving PIWIL3, the whole PIWIL3-TDRKH-PNLDC1 complex containing the bound piRNA could be immuno-precipitated with a TDRKH antibody.

Specifically, we used lysates of bovine oocytes as starting material to isolate endogenous PIWIL3-TDRKH-PNLDC1 complexes, and extracted the RNAs with Trizol to isolate small RNAs for sequencing. In parallel, we performed the same procedures on the input samples (oocytes lysate before IP), for comparison of piRNAs enrichment. The sequenced libraries contained about 30 million raw reads for input samples and IP samples. After trimming, length filtering (18-36 nt) and quality processing, approximately 13 million reads per replicate from TDRKH IP samples were mapped to the *Bos taurus* genome. In both duplicates of TDRKH immuno-precipitated piRNAs and total piRNAs (Input), high R^2^ correlations of 0.97-0.98 were obtained between biological (independent IP and sequencing) replicates (Figure S2a). The raw data for RNA sequencing is appended in Supplementary information, Table S3.

Next, we examined the RNA species in the TDRKH IP samples to characterize the small RNA pool associated with the PIWIL3/TDRKH complex. While miRNAs, rRNAs, and tRNAs were underrepresented in the TDRKH IP samples compared to input samples, piRNAs had clearly higher affinity for PIWIL3-TDRKH-PNLDC1 complexes (Figure 5a). Both input (70 %) and TDRKH IP (80 %) samples predominantly comprised small RNAs that map to annotated piRNA producing loci^20^ and exhibit a strong bias for uracil (U) at the 5’ position. The 5’ U bias was even stronger in the IP samples (Figure 5b; Supplementary information, Table S3). To further confirm the genuine presence of piRNAs, we assessed the RNA length distribution from the IP samples, which revealed a normal distribution about the peak of 24-27 nucleotides, strongly indicative of piRNA species. In addition, we also observed a strong enrichment of read pairs with 10 nt 5’ overlap, which is again highly indicative of the presence of primary and secondary piRNAs in the IP samples. These piRNA properties indicate that both primary and secondary piRNAs are bound by the PIWIL3-TDRKH complexes in bovine oocytes (Figure 5b-d, Figure S2b; S3).

**Figure 5:**
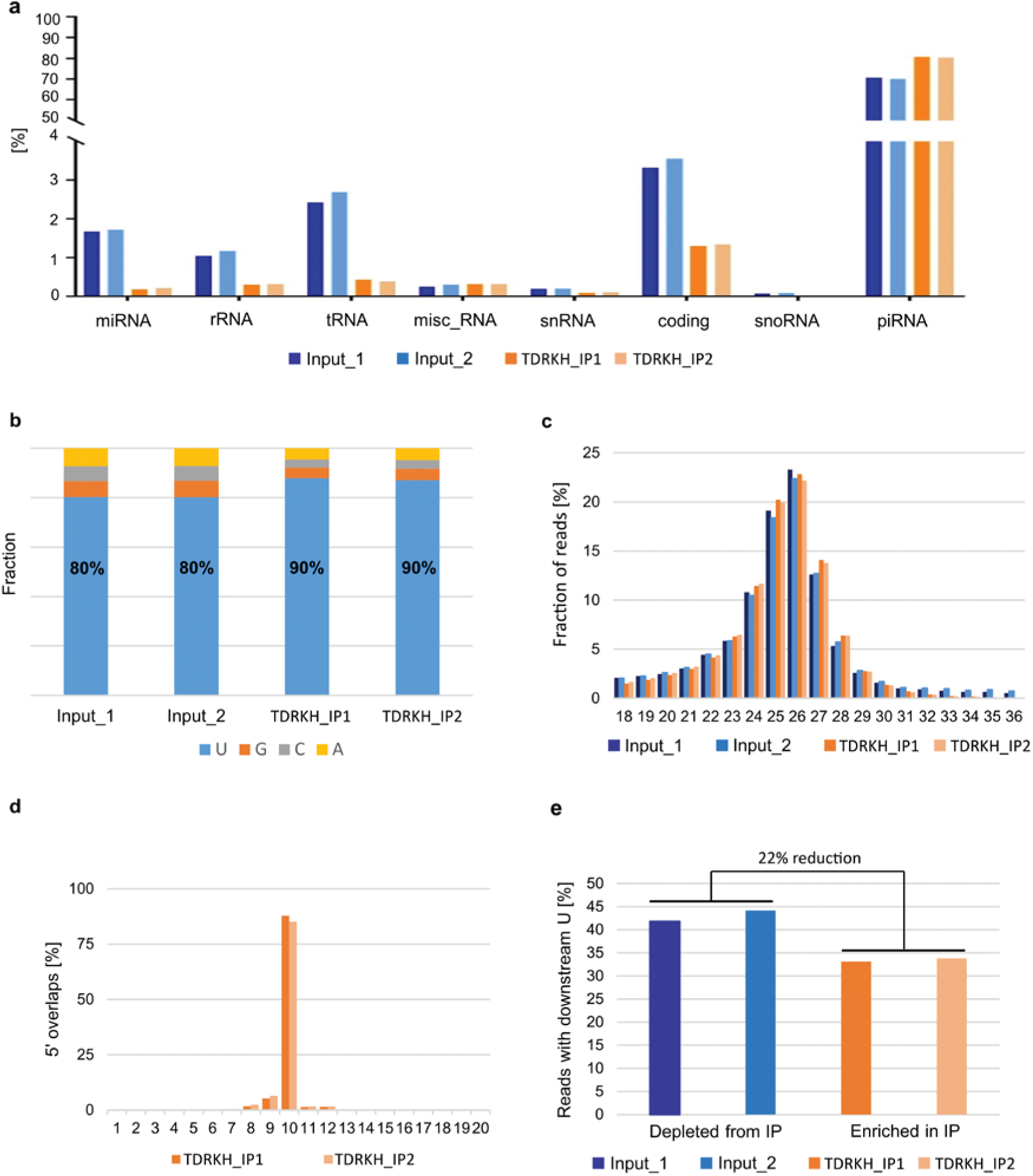
Analyses of piRNA sequences detected after TDRKH immunoprecipitation. Input 1, 2: duplicate total RNAs (blue). TDRKH_IP1, 2: duplicate immunoprecipitated RNAs (orange). **a** RNA categories detected in Input and TDRKH IP samples. **b** Nucleotide composition at the 1st position of the 5’ end. **c** Length distribution of the small RNA libraries. **d** Overlaps of 5’ends of reads that are mapped to opposite strands of the same locus. **e** Frequency of downstream 1U in the small RNAs. Percentage reduction in reads with downstream 1U calculated with respect to Input samples.

According to the current model of piRNA biogenesis, the endonuclease Zucchini (phospholipase D family member 6: PLD6) functions to promote piRNA biogenesis by cleaving pre-pre-piRNAs to generate the 5’ end of pre-piRNAs. A key characteristic of these intermediates is that they have a uridine directly downstream of the generated piRNA intermediate in the genome^21,22^. Once PIWI protein loaded with pre-piRNAs complexes with PNLDC1, 3’ end trimming by PNLDC1, a PARN family 3’-5’ exonuclease, is thought to start. Hence, piRNA intermediates on the TDRKH complex should lose their downstream U bias because of 3’ end trimming. In contrast, piRNA intermediates that are not loaded on the trimmer complex may still keep the downstream U bias. We evaluated this hypothesis based on our data, and indeed found a 22% reduction in downstream 1U with respect to Input samples (Figure 5e), indicative of trimming activity by the PIWIL3-TDRKH-PNLDC1 complex. Around 18% of the piRNAs in IP samples are marked by an extension of 1-2 nucleotides. Among it, piRNAs display adenylation predominantly, which is even more pronounced in the IP samples (Figure S5) indicating that TDRKH-IP piRNAs are more frequently trimmed and then adenylated.

Traditionally, piRNAs have been widely associated with TEs and repeat regions. To assess piRNAs in the mammalian PIWIL3/TDRKH complexes in relation to repetitive sequences, we compared the location of mapped small RNAs with the RepeatMasker annotation of the *Bos taurus* genome. We found that 45% of RNAs in TDRKH IP samples are mapped to transposon sequences, most notably LINE, SINE and LTR elements (Figure 6a). Within the LINE class, piRNAs map predominantly to L1 and BovB repeats, which is similar to the data previously reported for piRNAs extracted from regular oocyte samples (Figure S2c)^13,23^. Based on sequence homology a list was made of putative non-transposon target genes (Table S4). These genes are not only targeted but also processed via the ping-pong cycle, indicating that they are piRNA targets.

**Figure 6:**
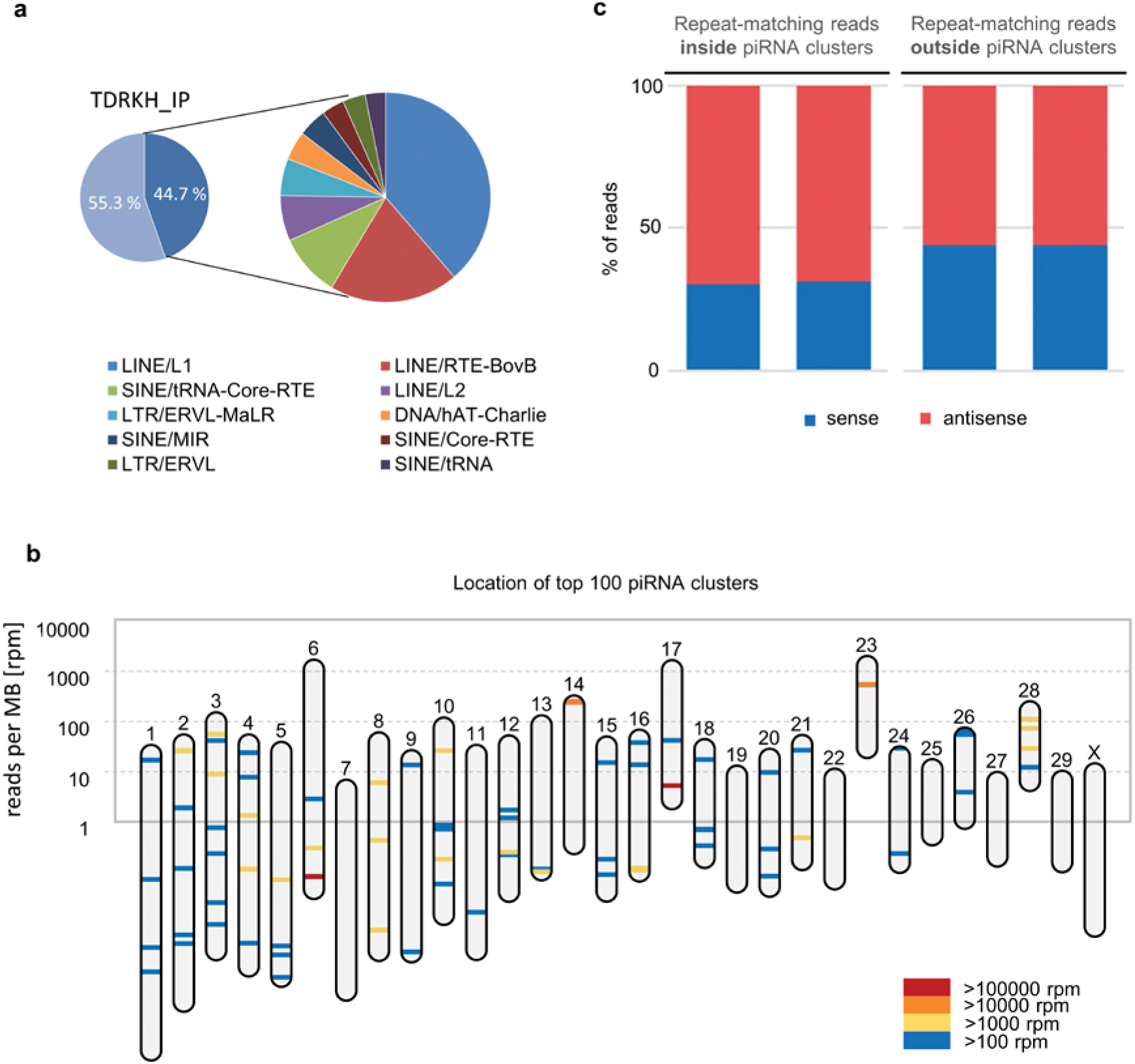
piRNAs mapping to transposons. **a** Pie chart depicting the transposon content (dark blue) of bovine piRNA populations from TDRKH IP samples, with further annotation in inset. **b** Mapping of piRNA clusters across chromosomes in the *Bos taurus* genome. Each piRNA cluster location is indicated by a bar colored according to its expression level. Y-axis refers to the number of clustered piRNAs per chromosome, normalized by chromosome size. **c** Repeat-matching reads inside and outside piRNA clusters in TDRKH IP samples.

We further addressed the chromosomal distribution of piRNA producing loci based on *de novo* piRNA cluster prediction with proTRAC (Figure 6b). We identified 674 distinct piRNA producing loci with a total size of 5 MB. In line with findings from other species, few piRNA clusters contribute to the majority of piRNA reads, as more than 80% of clustered piRNA reads were derived from only 12 predicted piRNA clusters up to 154 kb in length. Depending on the physical location of these piRNA clusters, the density of genomically encoded piRNAs per MB DNA can be diverse across different chromosomes. While the chromosomes 6, 17 and 23, which comprise the three largest piRNA clusters in terms of read counts, exhibit an overall density of >1600 aligned piRNA reads per MB, all other chromosomes exhibit on average of ∼50 aligned piRNA reads per MB. We further noticed that the identified piRNA clusters represent a main source of piRNAs antisense to transposon sequences, suggesting they are likely to be produced from piRNA clusters and regulate TEs. On average, 71% of transposon derived reads being reverse complementary to transposon sequences, while the same holds true for only 56% of transposon derived reads from outside of piRNA clusters (Figure 6c, Figure S2d).

## Discussion

Since a *PIWIL3* ortholog is absent from the mouse genome, the function of PIWIL3 has not been studied by gene knockout and has remained largely unknown. Based on genetic deletion of the other 3 Piwi genes *Miwi, Mili* and *Miwi2* in the mouse, it was hypothesized that in mammals PIWI proteins are only essential for male germ cells. Here we reveal on the contrary that PIWIL3 is expressed in the cytoplasm of bovine maturing oocytes. Our data demonstrate that in oocytes from antral follicles, pre-piRNAs/PIWIL3 associated with TDRKH and PNLDC1 on mitochondria, most likely revealing the piRNA biogenesis mechanism in mammalian oocytes.

TDRKH is a conserved Tudor domain protein localized on mitochondria that is critical for pre-piRNA 3’ end trimming in diverse animal models. As interacting partner of mouse MIWI, MIWI2 and a lesser extent to MILI, knockout of TDRKH resulted in spermatogenesis arrested in meiosis and reduced levels of mature piRNAs^16,24,25^. Various PIWI proteins have been suggested to interact with Tudor proteins through multiple arginine-glycine and arginine-alanine (RG/RA) rich clusters at their N termini in either an arginine methylation dependent, or independent manner depending on the species^16,17,26–28^. For the first time in a mammalian system, we identified a symmetrical glycine-arginine-alanine motif in the N terminus of PIWIL3 as a binding site for TDRKH in oocytes, and demonstrated indirectly that post-translational modifications on the side chain of R4 and R10 might be critical for TDRKH docking. Whether TDRKH binding requires the arginines to be methylated in PIWIL3 remains to be demonstrated, and indeed it has been shown that TDRKH preferentially recognizes unmethylated arginines in the N-terminus of PIWIL1^28^.

Interestingly, in bovine and human oocytes from antral follicles PIWIL3 was the only detected PIWI member^13,29^, while it was absent from testis tissue. TDRKH on the other hand was also strongly expressed in bovine testis and in mice is required for spermatogenesis indicating binding to a variety of Piwi molecules^16^. In mice, it has been reported that TDRKH acts as a key mitochondria-anchored scaffold protein that specifically recruits MIWI to the intermitochondrial cement and tethers PNLDC1 to couple MIWI recruitment and piRNA trimming during pachytene piRNA biogenesis. This TDRKH-mediated scaffolding function is essential for the production of MIWI-piRNAs, MIWI chromatoid body localization, transposon silencing and spermiogenesis^24^. We propose that in bovine oocytes, TDRKH is also important for PIWIL3 recruitment to mitochondria since PIWIL3 physically interacts with TDRKH and co-localizes with TDRKH on the mitochondria. This TDRKH binding brings the piRNA-bound PIWIL3 in close proximity to PNLDC1, a deadenylase responsible for the elimination of mRNA 3’ poly(A) tails and trimming during pre-pachytene and pachytene pre-piRNA maturation^25,30,31^.

In silkworms (*Bombyx mori*) it has been postulated that BmPapi (the silkworm homologue of TDRKH) recruits PIWI-pre-piRNA complexes on the surface of mitochondria to create an optimal platform for trimming. PNLDC1 binds with BmPapi and shortens the 3’ end of pre-piRNAs to the mature length after which the 3’ end is 2’-O-methylated by BmHen1^25^. In our data, sequences enriched in IP samples were slightly shorter (average size 25.2 nt of IP piRNAs versus 25.4 nt of input piRNAs). This was predominantly due to a smaller percentage of RNAs between 30 and 36 nt (Figure 5c). Moreover, with the support of reduced downstream U bias in the genome of IP samples, our results indicate that PNLDC1 is also responsible for the trimming of pre-piRNAs in mammalian oocytes. It is further suggested that these piRNAs are kept relatively unstable by post-transcriptional 3’-adenylation and are likely to be a signal for degradation possibly after activation of the embryonic genome^13^. Our qPCR data of expression of *PIWIL3, TDRKH* and *PNLDC1* mRNA indicate that the piRNA biogenesis complex in bovine embryos is mainly functional before the morula stage, also in agreement with a coordinated decrease in TDRKH and PNLDC1 levels during embryonic development detected previously^32^. While other PIWI proteins have been studied rigorously in lower model organisms, this study is the first to provide evidence in oocytes of mammals that piRNAs are present in a complex comprising both PIWIL3, TDRKH and PNLDC1.

PIWI proteins and piRNAs are regulators of both transposon and mRNA expression^33^. About 45% of the piRNA sequences detected in IP samples map to retrotransposon elements, while only fewer than 20% of piRNAs from adult bovine testis do so^20^, strongly indicating that PIWIL3 is involved in the control of retrotransposon elements in oocytes. piRNAs from oocytes and TDRKH-IPs map uniquely to intergenic regions of the genome, forming large piRNA clusters. While roughly half of the clustered piRNAs derive from transposon copies and are thus presumably involved in transposon control, the function of non-repeat derived piRNAs remain enigmatic. In agreement with Cuthbert et al^34^, PIWIL3-bound piRNAs clusters were predominantly detected on chromosomes 6, 17, 14 and 23. In contrast however, we observed enhanced piRNA mapping to chromosome 28, but not to chromosomes 8 and X. Since no other PIWI proteins were detected in bovine oocytes it seems unlikely that the differences are caused by oocyte piRNAs bound by other proteins, but more likely due to analysis settings.

During mouse spermatogenesis, meiosis induces a period of transcriptional quiescence which leads to the post-transcriptional silencing of transposon mRNAs by multiple epigenetic mechanisms that are in conjunction with the piRNA pathway^35^. However, since PIWIL3 is absent from mouse oocytes, piRNAs may play a more important role in oogenesis of non-murine mammals. We conducted a *PIWIL3* siRNA microinjection experiment with GV oocytes and oocytes matured to the MII stage. Despite huge efforts and efficient reduction in *PIWIL3* mRNA levels, the PIWIL3 protein level did not significantly change (data not show), which indicates that siRNA mediated knock down is not an efficient way to address the function of PIWIL3 in oocytes due to putatively long half-life of PIWIL3 proteins and limited culture time during oocyte maturation.

Some Argonaute family members including MIWI and human Agos are small RNA-guided nucleases (slicers) that catalyse cleavage of target nucleic acids depending on the presence of a conserved catalytic motif Asp-Glu-Asp-His inside the PIWI domain^36,37^. We also found the conserved catalytic motif DEDH in PIWIL3, leading us to speculate that PIWIL3 may act as a slicer at some point during oocyte meiosis and embryo development. This assumption is supported by the presence of a ping-pong signature, which indicates the cleavage of piRNA target transcripts, in simultaneous absence of other PIWI proteins. Further biochemical experiments should be performed to confirm this activity.

Depletion of *Piwi* genes lead to male-specify sterility because of defects in sperm formation. However, the functions of PIWI proteins in mammalian females are still largely unknown. For a long time, the function and piRNA-binding properties of PIWIL3 was not investigated, primarily due to the lack of animal systems expressing this important piRNA processing regulator, and the perennial shortage of PIWIL3-specific antibodies. Here we provide a first glimpse into the PIWIL3 piRNA repertoire in bovine oocytes. In future studies, mechanism and function of PIWIL3/TDRKH/PNLDC1 complexes in mammalian oocytes should be further investigated using knockout animal models, although the mRNA injection workflow we present here could also be useful for experiments with relatively shorter time durations.

## Methods

### Oocytes collection, maturation and fertilization

Bovine ovaries were collected from a local slaughterhouse and transported to the laboratory in a polystyrene box at room temperature (RT) within 2 h after slaughter. After washing and removal of extraneous connective tissue, the ovaries were transferred to a flask containing 0.9% NaCl supplemented with 10 U/ml penicillin/streptomycin (Gibco BRL, Paisley, UK) and maintained in a water bath at 30 °C. Cumulus oocyte complexes (COCs) were collected, matured and fertilized *in vitro* as described previously^38^.

### RNA isolation, cDNA synthesis

An RNeasy Micro Kit (Qiagen, Valencia, CA, USA) was used to extract total RNA as described before^39^. The columns were eluted with 18 μl RNAse-free water. For cDNA synthesis, 10 μl RNA was mixed with 4 μl 5 x RT buffer (Invitrogen, Breda, The Netherlands), 0.2 μl RNAsin (Promega, Leiden, The Netherlands), 0.75 μl Superscript III reverse transcriptase (Invitrogen), 0.4 μl random primers (Invitrogen), 2 μl DTT (Invitrogen) and 1 μl dNTP (Promega). Minus RT blanks were prepared from 10 μl of the same RNA sample under the same conditions without the addition of reverse transcriptase. The mixture was incubated for 1 h at 55 °C, followed by 5 min at 80 °C before storage at –20 °C.

### PIWIL3 plasmids construction

PIWIL3 cDNA was amplified from oocyte cDNA using PCR. PIWIL3 plasmid construction primers are shown in supplementary table S2a. The DNA was cloned into pcDNA3-EGFP, a gift from Doug Golenbock (Addgene plasmid # 13031). A Q5® site-directed mutagenesis kit (New England BioLabs, Ipswich, MA, USA) with forward primer 5’-CTACAGAAGAAAGCACAATG −3’ and reverse primer 5’-AAGGTAAAAGAGAGATTTTGAC −3’ was used to delete the stop codon in *PIWIL3*. The HiScribe™ T7 ARCA mRNA kit (New England BioLabs) was used to transcribe *PIWIL3-EGFP, EGFP-PIWIL3* and *EGFP* mRNA *in vitro*. The mRNA integrity was checked by electrophoresis on 1% agarose gels with MOPS buffer prior to microinjection.

### Injection of mRNA into oocytes

Before injection, *EGPF-PIWIL3, PIWIL3-EGFP* and *EGFP* mRNA were diluted with RNase-free water to a final concentration of 100 nM. Subsequently, 6 denuded germinal vesicle (GV) stage oocytes were transferred to a 5 μl drop of HEPES buffered M199 with 10% FCS in a 60 mm dish overlaid by mineral oil at 37 °C on an IX71 inverted microscope (Olympus, Leiderdorp, the Netherlands) equipped with a heated stage at 38.5 °C. In total, 5 μl of the mRNA was loaded into a microinjection needle with a 30° angle and 4.3-4.9 μm inner diameter of the tip (Origio, Vreeland, The Netherlands). Injection was performed at 100 hpa for 0.2 seconds. After injection, oocytes were cultured in maturation medium with 25 μm Roscovitine for 8 h to collect GV stage oocytes; only oocytes with green fluorescence were washed in PBS and fixed in 4% paraformaldehyde (PFA) 30 min at room temperature (RT). Zygotes used for microinjection were collected 8 h after in vitro fertilization (IVF).

### Immunoblotting

Groups of 60-100 injected oocytes or 100 normal oocytes, ovary and testis tissues were lysed in RIPA buffer (Pierce Biotechnology, Rockford, IL, USA) supplemented with 1% protease/phosphatase inhibitor (ThermoFisher, Waltham, MA, USA). Lysates were separated by electrophoresis on 8% SDS-PAGE gels and subsequently transferred to nitrocellulose membranes (Bio-Rad, Hercules, CA, USA). Blots were blocked with 5% milk in TBST (TBS + 0.1% Tween 20) for 1 h at RT and incubated with GFP antibody (1:1000, sc-9996, Santa Cruz Biotechnology, Dallas, TX, USA) or TDRKH antibody (1:1000, 13528-1-AP, Proteintech) overnight at 4 °C. Blots was washed three times (10 min each) in PBST followed by 1 h incubation with secondary antibody: HRP-conjugated goat anti mouse IgG (1:5000, sc-2005, Santa Cruz Biotechnology) or HRP-conjugated goat anti rabbit IgG (1:5000, 31460, Pierce Biotechnology) at RT. Antibody binding was detected using ECL Super Signal West Dura Extended Duration Substrate (ThermoFisher) and exposure to maxAgfa CL-XPosurelight films (ThermoFisher).

### Mitochondrial staining and immunofluorescence of bovine oocytes

For mitochondrial staining, oocytes and early stage embryos were incubated in M199 with 500nM MitoTracker™ Red CMXRos (M7512, ThermoFisher) for 1 h in a humidified incubator at 38.5 °C and 5% CO_2_. Oocytes and embryos were subsequently washed 3 times in PBS and fixed in 4% PFA for 30 min at RT.

Immunofluorescence was conducted largely as described before^39^. After fixation, oocytes were washed three times in PBST (PBS+10% FBS+0.1% Triton-100), permeabilized for 30 min using 0.5% Triton-X100 in PBS with 10% FBS and blocked in PBST for 1 h at RT. Incubation with TDRKH antibody (13528-1-AP, Proteintech ThermoFisher) 1:100 was at 4°C overnight. Oocytes were then washed three times in PBST for 15 min each and incubated with secondary goat anti-rabbit 1:100 (AlexaFluor 488, Life Technologies, Bleiswijk, the Netherlands) for 1 h in the dark at RT. After washing, oocytes were incubated with 4’,6-diamidino-2-phenylindole (DAPI) for 20 min and mounted onto glass slides using Vectashield (Vector Laboratories, Burlingame, CA, USA). Fluorescence was examined by confocal laser scanning microscopy (TCS SPE II, Leica, Wetzlar, Germany).

### Immunohistochemistry

Tissues were fixed in 4% PFA overnight at 4 °C and embedded in paraffin. Sections (5 µm) were deparaffinised, washed in water and citrate buffer (pH 6.0) for antigen retrieval followed by blocking with BSA. The sections were treated with the primary anti-TDRKH antibody (1:200, 13528-1-AP, Proteintech ThermoFisher) overnight at 4 °C. After incubation with HRP-conjugated goat anti rabbit secondary antibody (1:5000, #31460, Pierce Biotechnology), the slides were washed and developed with Diaminobenzidine (DAB) solution, K3468, Dako) and counterstained with hematoxylin. As a negative control, sections were incubated with rabbit Immunoglobulin (X0903, Dako Agilent, Santa Clara, CA, USA).

### Immunoprecipitation and MS identification

Cells were lysed in IP lysis buffer (Pierce™ Crosslink IP Kit, 26147, ThermoFisher) and 10 µg TDRKH antibody (13528-1-AP, Proteintech) or rabbit Immunoglobulin (X0903, Dako Agilent) was coupled to Protein A/G-coated agarose beads, and crosslinked with 2.5 mM DSS to prevent co-elution. The antibody conjugated beads were mixed with bovine oocyte lysate incubated for 60 min to retrieve TDRKH complexes. The IP eluates were reduced with 4 mM DTT, alkylated with 8mM Iodoacetamide, and then digested with 1:75 trypsin at 37 °C overnight. The resulting peptides were desalted by c18, dried by vacuum centrifugation and reconstituted in 10% formic acid for analysis on an Orbitrap Q Exactive HF spectrometer (ThermoFisher) connected to a UHPLC 1290 LC system (Agilent). Raw files were processed using Proteome Discoverer 1.4.1.14 and searched using Mascot against the bovine database (downloaded on 2017.11.29, containing 32206 entries). Cysteine carbamidomethylation was used as a static modification and methionine oxidation was set as a possible dynamic modification. Up to 2 missed cleavages were allowed. False discovery rates of 1% was set for both protein and peptide identification. For IP samples prepared for Western blot, 1 mM DSSO was used to crosslink protein complex before elution.

### Quantitative RT-PCR

RNA isolation and cDNA synthesis were described as before in the methods. 20-30 oocytes or embryos were collected for each sample. Minus RT blanks were prepared from 5 μl of the same RNA sample under the same conditions, but without addition of reverse transcriptase. 1 μl of the resulting 20 μl cDNA was used for qRT-PCR analyses with the primers listed in Supplementary Table S2b. The qRT-PCR reactions were performed using a real-time PCR detection system (MyiQ Single-color Real-Time PCR Detection System; Bio-Rad Laboratories, Hercules, CA, USA) with IQ Sybr Green Supermix (Bio-Rad Laboratories). Three biological repeats were analyzed. The relative starting quantity for each experimental sample was calculated based on the standard curve made for each primer pair. Data normalization was performed using *GAPDH* and *SDHA* as reference genes with the same set of samples.

### IP for piRNA sequencing

Bovine GV stage oocytes were isolated and snap frozen in liquid nitrogen. Upon collection, 500 oocytes per IP sample were taken up in 500 µL cold lysis/IP buffer (25 mM Tris pH 7.5, 150 mM NaCl, 1.5 mM MgCl2, 1% Triton-X100, 1 mM DTT, Protease Inhibitor (04693159001, Roche)) and sonicated 3 x 30 sec. The samples were centrifuged at 12 000 g at 4 ^o^C and the supernatant was used for IP. Samples were incubated with anti-TDRKH 1:100 (13528-1-AP, Proteintech ThermoFisher) for 2 h at 4 °C, rotating. Next, samples were incubated for another 45 min at 4 °C with 30µL pre-washed Dynabeads protein G. Afterwards, the beads were washed 3 times with cold wash buffer (25 mM Tris pH 7.5, 300 mM NaCl, 1.5 mM MgCL2, 1 mM DTT) and taken up in Trizol (15596018, ThermoFisher), followed by RNA and protein isolation according to the manufacturer. The RNA samples were used for library preparation.

### Library preparation

NGS library prep was performed with NEXTflex Small RNA-Seq Kit V3 following Step A to Step G of Bioo Scientific’s standard protocol (V16.06). Libraries were prepared with a starting amount of 5ng and amplified in 25 PCR cycles. Amplified libraries were size selected on a 8% TBE gel for the 18 – 40 nt insert size fraction. Libraries were profiled in High Sensitivity DNA Assay on 2100 Bioanalyzer (Agilent) and quantified using the Qubit dsDNA HS Assay Kit, in a Qubit 2.0 Fluorometer (Life Technologies). All 4 samples were pooled in equiAL molar ratio and sequenced on 1 NextSeq 500 Flowcell, Highoutput 75-cycle-kit, SR for 1x 84 cycles plus 7 cycles for the index read.

### Sequencing data processing

Trimming of adapter sequences comprising flanking 4 nucleotide random tags with subsequent filtering of putative PCR duplicates was performed as previously described^13,40^. Trimmed sequence reads <18 or >36 nt in length as well as low complexity reads were filtered and the remaining reads were collapsed to non-identical sequences using Perl scripts from the NGS TOOLBOX^13^. Mapping to the genome of *Bos taurus* (bosTau7) and subsequent filtering of alignments was performed as described^41^. The obtained map files were used as input for unitas small RNA annotation pipeline (v.1.5.3, additional options: -pp –skip_dust)^42^. Reference sequences were obtained from Ensembl Database (release 93)^43^, GtRNAdb^44^, piRNA cluster database^20^, SILVA rRNA database (release 132) and miRBase (release 22)^45,46^. The amount of sequences that correspond to repetitive DNA was determined based on RepeatMasker annotation (RepeatMasker open-4.0.5, Repeat Library 20140131) using the custom Perl script RMvsMAP.pl. *De novo* piRNA cluster prediction was performed with proTRAC 2.4.2 for each dataset separately using default settings. The predicted cluster coordinates were merged as described^41^. piRNA target genes were predicted as described before without allowing any mismatches when mapping piRNAs to coding sequences^47^.

## Acknowledgements

We are grateful to Prof. René Ketting (Institute of Molecular Biology, Mainz) for providing access to the small RNA sequencing facility. We also thank Dr. Pieter van Breugel for assistance in plasmid construction. Minjie Tan is supported by a PhD scholarship from the China Scholarship Council.

## Author contributions

M.T, W.W and B.R designed the study and interpreted the data. M.T collected primary materials and performed plasmid construction, microinjection, and immunostaining. M.T and M.J.D performed IP experiments and MS analyses. H.T performed qRT-PCR experiments. E.F.R performed RNA sequencing. D.R performed the small RNA sequencing analysis. M.T, B.R, W.W and T.S. wrote the manuscript. All authors contributed and approved the final version of the manuscript.

## Material and Correspondence

All data and material that support the findings of this study are available from the corresponding authors Wei Wu or Bernard A.J. Roelen upon reasonable request.

## Supplementary Information

**Fig. S1:**
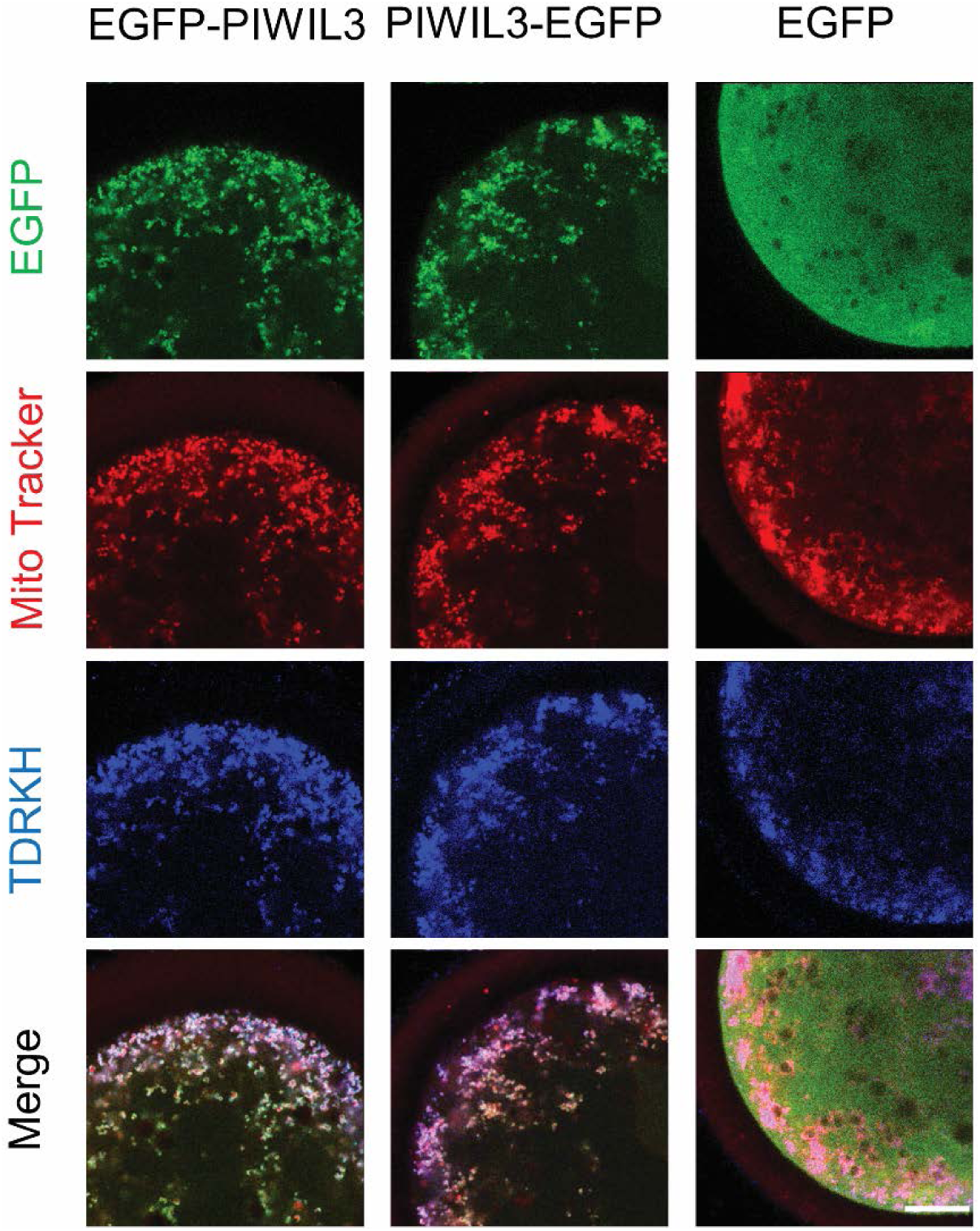
Fluorescent localisation after microinjection of PIWIL3-EGFP or EGFP-PIWIL3 mRNA into GV stage bovine oocytes. Distribution of EGFP-PIWIL3 and PIWIL3-EGFP (green), MitoTracker (mitochondria, red) and TDRKH (blue) in oocytes. Scale bar, 20 μm.

**Figure S2:**
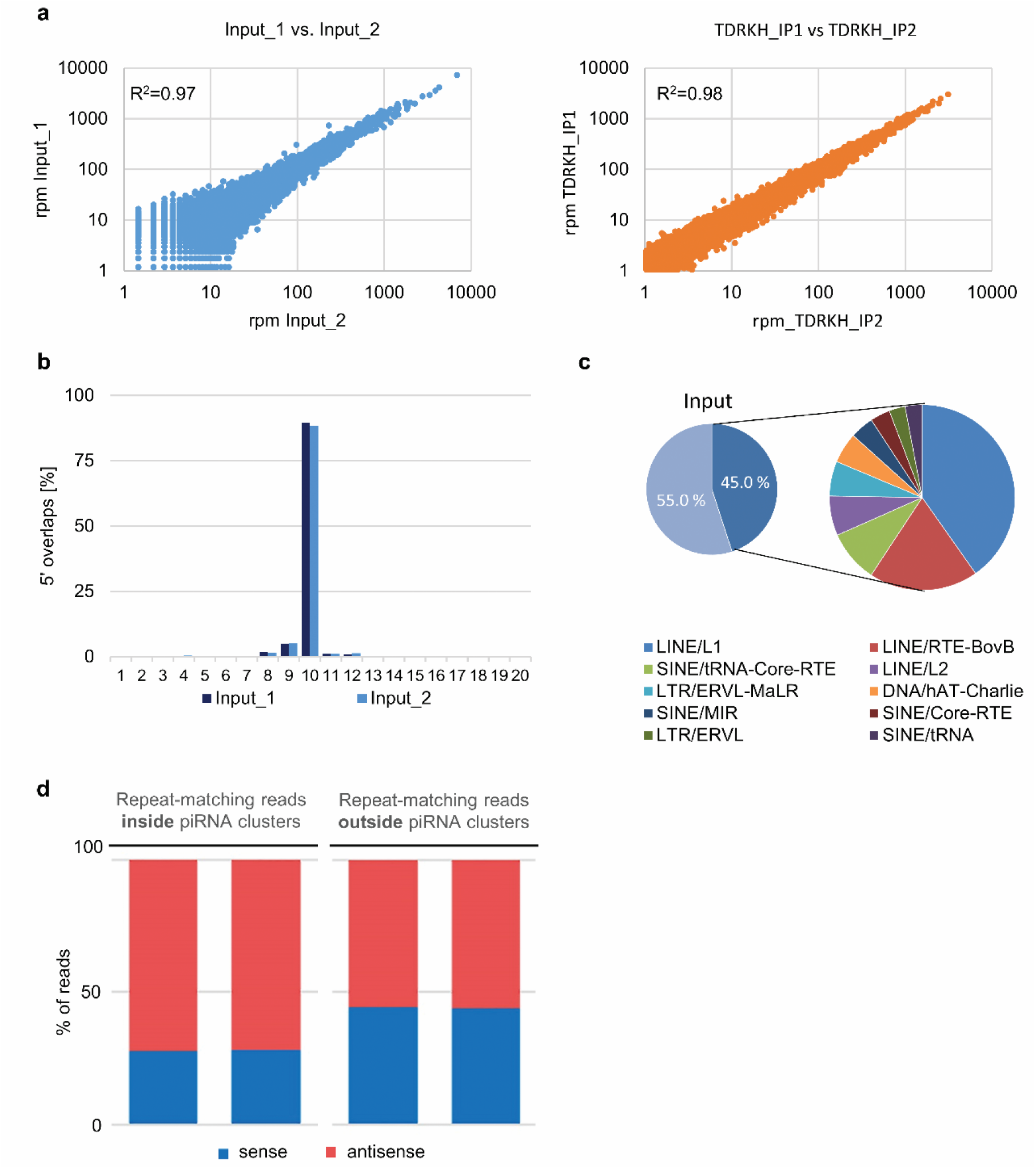
piRNA sequencing summary statistics. **a** Technical reproducibility. Pearson correlation between sequencing replicates varied in the range of 0.97<r^2^<0.98. rpm, reads per million. **b** Overlaps of 5’ ends of reads that are mapped to opposite strands of the same locus. **c** Pie chart depicting the transposon content of bovine piRNA populations from Input samples. **d** Repeat-matching reads inside and outside piRNA clusters for Input samples.

**Figure S3:**
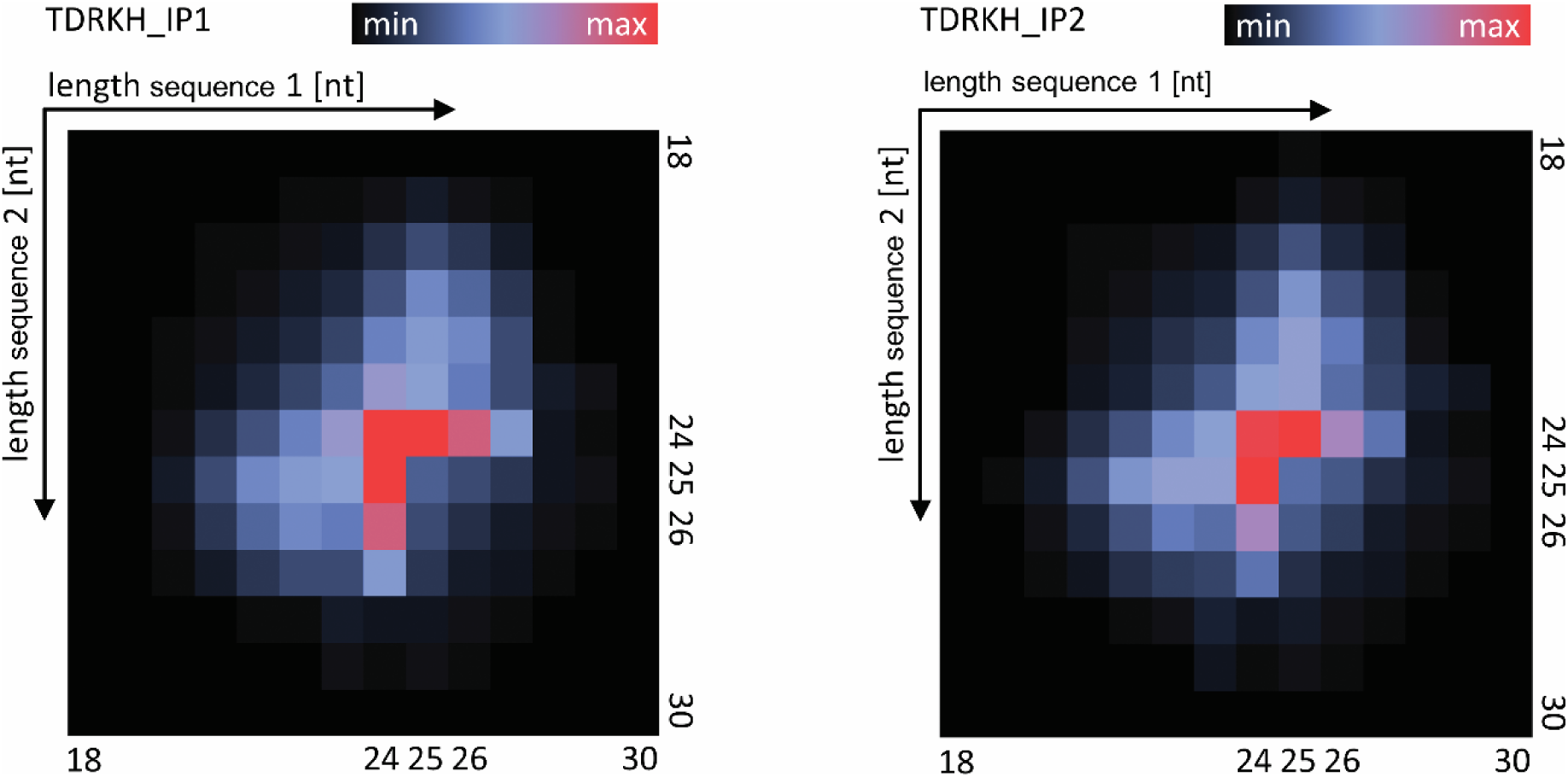
Sequence length analysis of piRNAs that participate in the ping-pong amplification loop. Ping-pong matrices illustrate frequent length-combinations of ping-pong pairs (sequences with 10 bp 5’ overlap), indicated in red. X-axis and Y-axis refer to sequence read length of the two sequences of a ping-pong pair.

**Figure S4:**
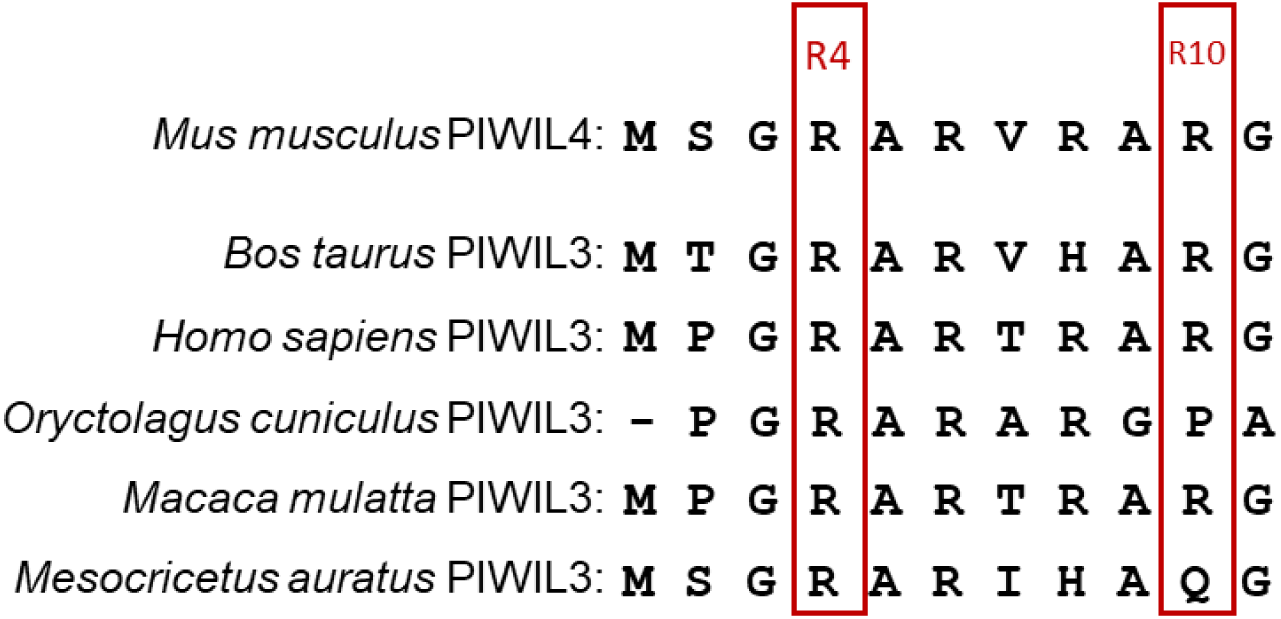
Sequence alignment of mammalian PIWI N terminal domain. R4 and R10 illustrate the arginines in *Bos taurus* PIWIL3 mutated in our experiment.

**Figure S5:**
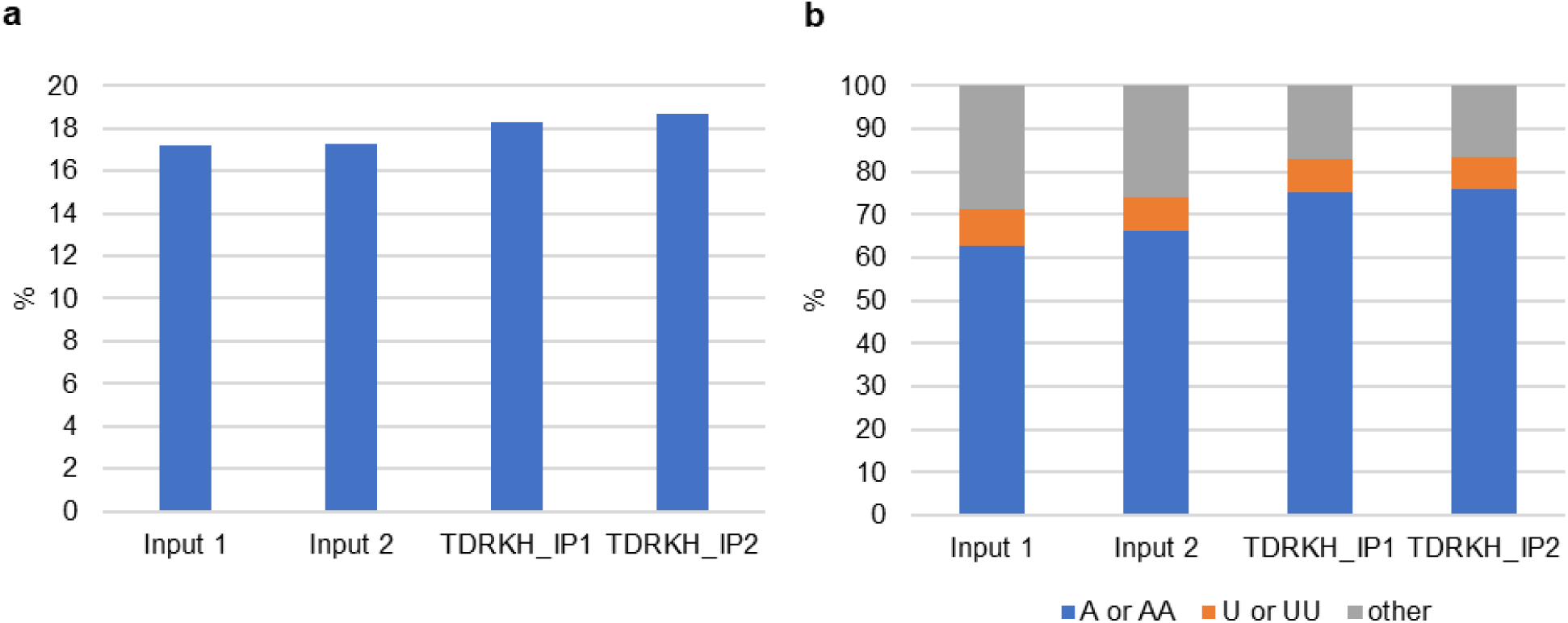
Non-templated nucleotide analysis. **a** Frequencies of non-templated nucleotides at the 3’ end from the indicated libraries. **b** Frequencies of the identified non-templated nucleotides into “A”, “U” or “other” tails.

**Table S2:**
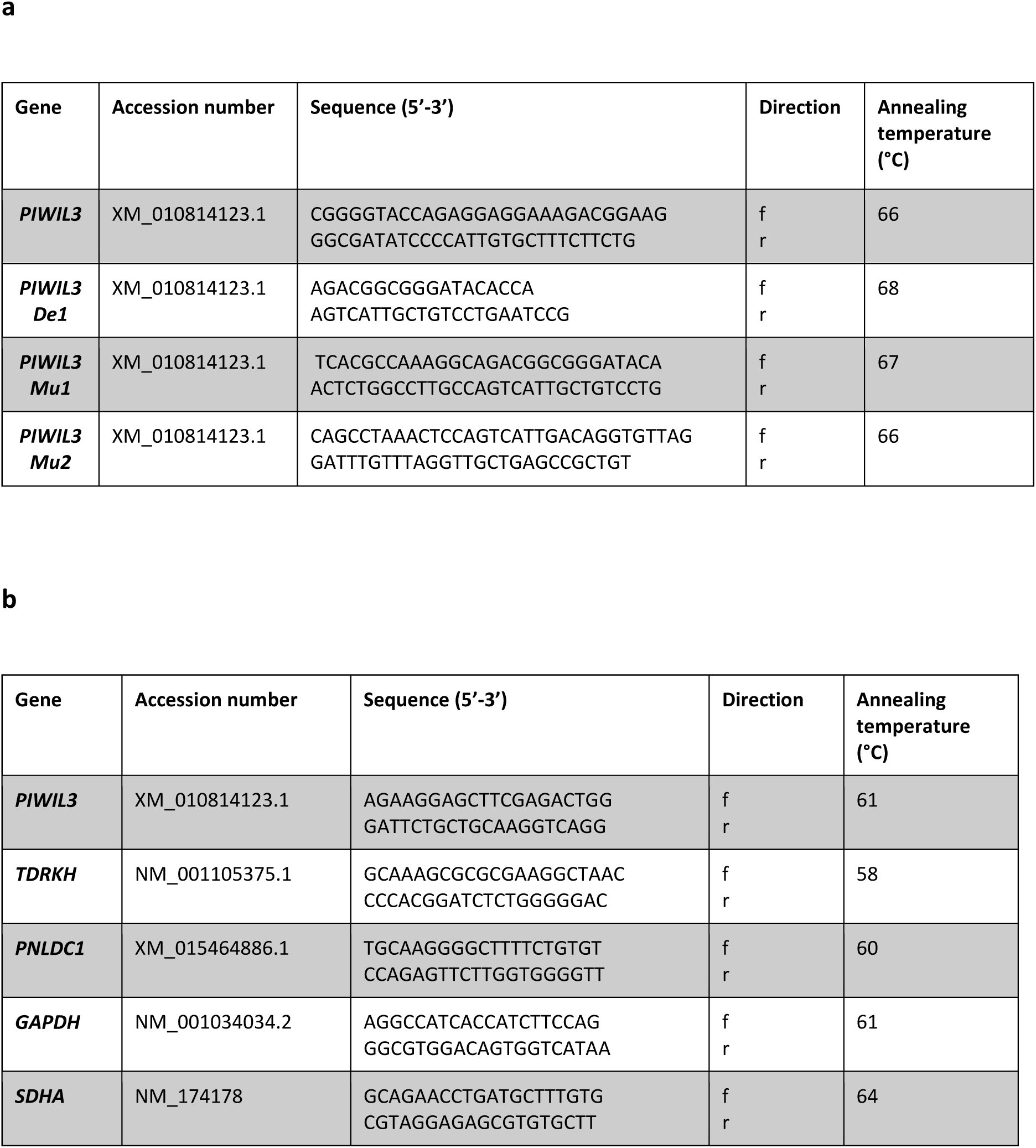
Primers for PIWIL3 (a) plasmid construction and (b) qRT-PCR. “f” and “r” indicate forward and reverse respectively.

**Table S1 TDRKH IP MS data (Excel file)**.

**Table S3: Small RNA sequencing data (Excel file)**.

**Table S4: Non-transposon putative target transcript genes (Excel file)**.

